# miR-206 knockout shows it is critical for myogenesis and directly regulates newly identified target mRNAs

**DOI:** 10.1101/2020.01.03.894246

**Authors:** Georgiana M. Salant, Kimngan L. Tat, James A. Goodrich, Jennifer F. Kugel

## Abstract

The muscle specific miRNA, miR-206, is important for the process of myogenesis; however, studying the function of miR-206 in muscle development and differentiation still proves challenging because the complement of mRNA targets it regulates remains undefined. In addition, miR-206 shares close sequence similarity to miR-1, another muscle specific miRNA, making it hard to study the impact of miR-206 alone in cell culture models. Here we used CRISPR/Cas9 technology to knockout miR-206 in C2C12 muscle cells. We show that knocking out miR-206 significantly impairs and delays differentiation and myotube formation, revealing that miR-206 alone is important for myogenesis. In addition, we use an experimental affinity purification technique to identify new mRNA targets of miR-206 in C2C12 cells. We identified over one hundred mRNAs as putative miR-206 targets. Functional experiments on six putative targets indicate that Adam19, Bgn, Cbx5, Smarce1, and Spg20 are direct miR-206 targets in C2C12 cells. Our data show a unique and important role for miR-206 in myogenesis.

## Introduction

MicroRNAs (miRNAs) are small ~22 nucleotide (nt) non-coding RNAs that play pivotal roles in controlling gene expression (1). They negatively regulate gene expression by binding to target mRNAs to trigger degradation of the mRNA or to repress translation of the mRNA into protein (1). Most often miRNAs bind to the 3’UTR of their mRNA target using a seed sequence, a conserved 6-8 nt sequence on the 5’ end of the miRNA that base pairs perfectly with the target mRNA (2). A single miRNA has the potential to target many different mRNAs, hence post-transcriptional repression by a miRNA can have significant downstream effects on regulatory pathways in cells. Indeed, miRNAs have been shown to regulate many different biological processes and cellular fate decisions, including proliferation, differentiation, apoptosis, and tissue development (3). In addition, misregulation of miRNAs contributes to diseases such as cancer and skeletal muscle disorders (4–6).

One biological program in which miRNAs play an important regulatory role is myogenesis, the process of generating skeletal muscle (7). Myogenesis occurs during embryonic development as well as in adult skeletal muscle cells after injury (8). Several muscle specific miRNAs, termed myomiRs, are integral for the myogenic programming to occur (9, 10). MyomiRs facilitate myogenesis by both promoting muscle differentiation and inhibiting proliferation and/or alternative cell fates (5, 10). Moreover, myomiRs are thought to be important in skeletal muscle diseases, although the mechanisms by which they function in promoting and maintaining disease states remain ill-defined (5, 10). Among the myomiRs, miR-1, miR-133, and miR-206 are the most widely studied. MiR-1 and miR-133 are specifically expressed in both cardiac and skeletal muscle, whereas miR-206 is exclusively expressed in skeletal muscle (5, 10). The expression of each of these myomiRs is highly upregulated during embryogenesis and during differentiation of myoblasts to myotubes in cell culture models of myogenesis (11–17).

MiR-206 is the focus of the studies described here. Previous work provides evidence that miR-206 is critical for myoblast differentiation. In cell culture models, transient knockdown of miR-206/miR-1 with miR-133 disrupts proper myotube formation (12, 18). Moreover, a mouse knockout of miR-206 shows decreased satellite cell differentiation and a delay in myofiber growth after cardiotoxin injury, suggesting a role in skeletal muscle regeneration (19). MiR-206 also protects against neuromuscular disease progression since DMD (Duchenne muscular dystrophy) and ALS (amyotrophic lateral sclerosis) mouse models that lack miR-206 demonstrate accelerated muscle dysfunction (19, 20). Although it is clear that miR-206 has a role in muscle cell development, the molecular mechanisms by which it functions are more poorly defined, in part because the complete set of mRNAs targeted by miR-206 is not known.

A handful of mRNA targets of miR-206 have been experimentally identified. Their downregulation aids in skeletal muscle differentiation by impacting diverse biological pathways. For example, miR-206 impacts DNA synthesis and cell cycle progression by downregulating DNA polymerase alpha subunit 1 (Polα1) (12) and Cyclin D1 (Ccnd1) (21), respectively; controls cell communication and cytoskeletal structure by downregulating gap-junction protein alpha 1 (Gja1) (22) and utrophin (Utrn) (23), respectively; and affects RNA splicing by targeting serine and arginine rich splicing factor 9 (Srsf9) (24). Hundreds of additional target mRNAs are predicted for miR-206 using computational approaches; however, the fact that miRNAs require only limited complementarity to bind an mRNA target makes computational-based targetprediction difficult. Furthermore, miR-206 shares an identical seed sequence with miR-1 and the fully processed 22 nt miRNAs differ in sequence at only 4 positions (5, 25). Consequently, both miRNAs likely target many of the same mRNAs. Indeed, experimental approaches to knockdown miR-206 or miR-1 with antagomiRs, for example, result in both being knocked down (18, 26).

Here we studied the role of miR-206 during skeletal muscle differentiation using mouse C2C12 cells as a model system for myogenesis. We created a knockout (KO) of the miR-206 gene in C2C12 cells using CRISPR/Cas9 technology, which eliminated the miR-206 without affecting miR-1 levels. We found that in the KO cells, the absence of miR-206 significantly delayed myotube formation, indicating a unique role for miR-206 in myogenesis that cannot be compensated for by miR-1. To identify new candidate mRNA targets of miR-206, we purified miR-206-containing complexes from C2C12 cells before and after differentiation and sequenced the associated mRNAs. We identified 130 putative miR-206 targets and selected six to experimentally validate in the WT and KO cells. The majority of those tested show promise as bona fide targets for regulation by miR-206. Together our data show a unique and important role for miR-206 in myotube formation and identify novel pathways by which this miRNA might elicit its effects.

## Results

### Knockout of miR-206 in C2C12 cells causes delayed differentiation of myoblasts to myotubes

To explore the specific role of miR-206 in myogenic differentiation, we used CRISPR/Cas9 technology to generate a miR-206 knockout (KO) C2C12 cell line. We compared miR-206 expression levels before and after differentiation in the KO cell line versus wild-type (WT) C2C12 cells using RT-qPCR (Figure 1A). A dramatic increase (~50-fold) in miR-206 expression was observed in WT cells after four days of differentiation, which was expected (27). By contrast, miR-206 was not detected above background in the KO cell line before or after growth in differentiation media, confirming perturbation of the miR-206 locus.

**Figure 1.**
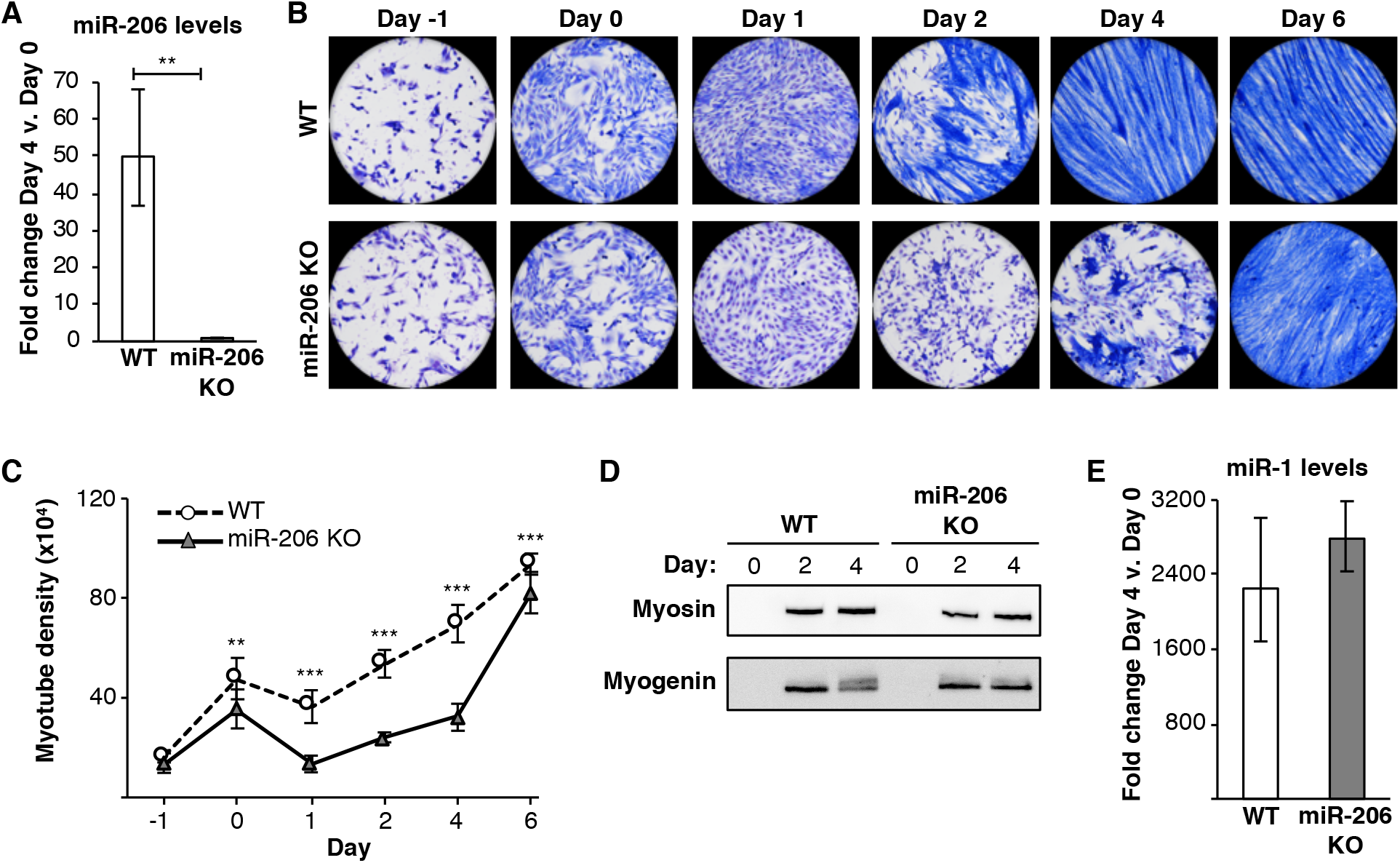
A miR-206 knockout (KO) cell line shows a delay in myotube formation upon differentiation. (A) miR-206 expression does not increase in miR-206 KO cells incubated in differentiation media. Average fold change in miR-206 expression, normalized to control sno202 RNA levels, on Day 4 relative to Day 0 is plotted. The error bars denote the standard deviations (n=3). ** indicates a p-value < 0.01, as determined from an unpaired two-tailed t-test. (B) miR-206 KO cells show a delay in myotube formation upon treatment with differentiation media compared to WT cells. Representative images of Jenner-Giemsa stained C2C12 WT and miR-206 KO cells during a time course of treatment with differentiation media are shown. Cells were switched to differentiation media on Day 0. (C) miR-206 KO cells show significantly less myotube density upon treatment with differentiation media compared to WT cells. Cells were switched to differentiation media on Day 0. Myotube density was calculated as the sum of pixels attributed to tones 0-75 at each time point (28). The data are the average of twelve imaged regions from three biological replicates, and the error bars denote the standard deviations (n=12). ** indicates a p-value < 0.01 and *** indicates a p-value < 0.001, as determined from an unpaired two-tailed t-test. (D) Myosin and myogenin proteins are expressed at similar levels in WT and miR-206 KO cells. Shown are Western before and after treatment with differentiation media. (E) miR-1 expression is induced to similar levels in WT and miR-206 KO cells after culturing in differentiation media. Average fold change in miR-1 expression, normalized to control sno202 RNA levels, on Day 4 relative to Day 0 is plotted; the error bars denote the standard deviation (n=3).

To investigate the effect of miR-206 KO on cellular phenotypes and myotube formation, we stained cells with Jenner-Giemsa dyes, a dual histological staining method that stains myoblasts light pinkish-purple and myotubes dark blue (28). WT and KO cells were stained at multiple time points over 6 days of growth in differentiation media (Figure 1B). Myotubes formed in WT cells by Day 2, as shown by the thick, multinucleated, dark blue cells. Moreover, myotubes continued to grow and align through Day 4 and persisted into Day 6. By contrast, miR-206 KO cells lacked myotube formation at Day 2, as evident by minimal dark blue staining and no elongated, multinucleated cells. By Day 4 dark blue staining was present, marking the start of myotube formation; however, these myotubes were stunted and lacked proper multinucleation and elongation compared to WT cells. Even by Day 6 where the entire field of view showed dark blue staining, the myotubes appeared stringy and lacked the thickness observed in WT cells, pointing to a persistent deficiency in myoblast fusion.

We quantified myotube formation in WT and miR-206 KO cells over the 6-day time course. To do so, WT and KO cells were imaged in grayscale and the pixel tones that corresponded to myotubes were summed. As plotted in Figure 1C, a significant difference in myotube density between WT and miR-206 KO cells was observed throughout the time course in differentiation media. We also determined the effect of miR206 knockout on levels of two canonical protein markers of muscle cell differentiation, myogenin and myosin. As shown in the immunoblots in Figure 1D, both of these proteins were expressed at similar levels in the WT and KO cells after growth in differentiation media, despite the delayed differentiation phenotype and failed elongation of myotubes in the miR-206 KO cells. Hence, knockout of miR206 is uncoupled from the expression level of these two differentiation markers. We tested whether the lack of miR-206 had a secondary effect on miR-1 levels during differentiation. We found that miR-1 levels increased to a similar extent in the two cell lines after four days in differentiation media (Figure 1E). Taken together, these data show that miR-206 has an important and non-redundant role with miR-1 in myogenic differentiation.

### Known targets of miR-206 are differentially regulated in miR-206 KO and WT cells

We next determined how knock out of miR-206 affected known mRNA targets. We investigated Ccnd1 and Gja1, two experimentally validated miR-206 mRNA targets that are downregulated upon differentiation in C2C12 cells (21, 22). We measured the fold change in levels of the two mRNAs after four days in differentiation media in both the miR-206 KO and WT cell lines using RT-qPCR (Figure 2A). In WT cells (white bars), levels of Ccnd1 and Gja1 significantly decreased upon differentiation, as expected. In miR-206 KO cells (gray bars), we also saw a significant decrease in the levels of these two mRNAs, albeit slightly less than in the WT cells. This indicates that that the differentiation-induced downregulation of these mRNAs is only partially dependent on miR-206.

**Figure 2.**
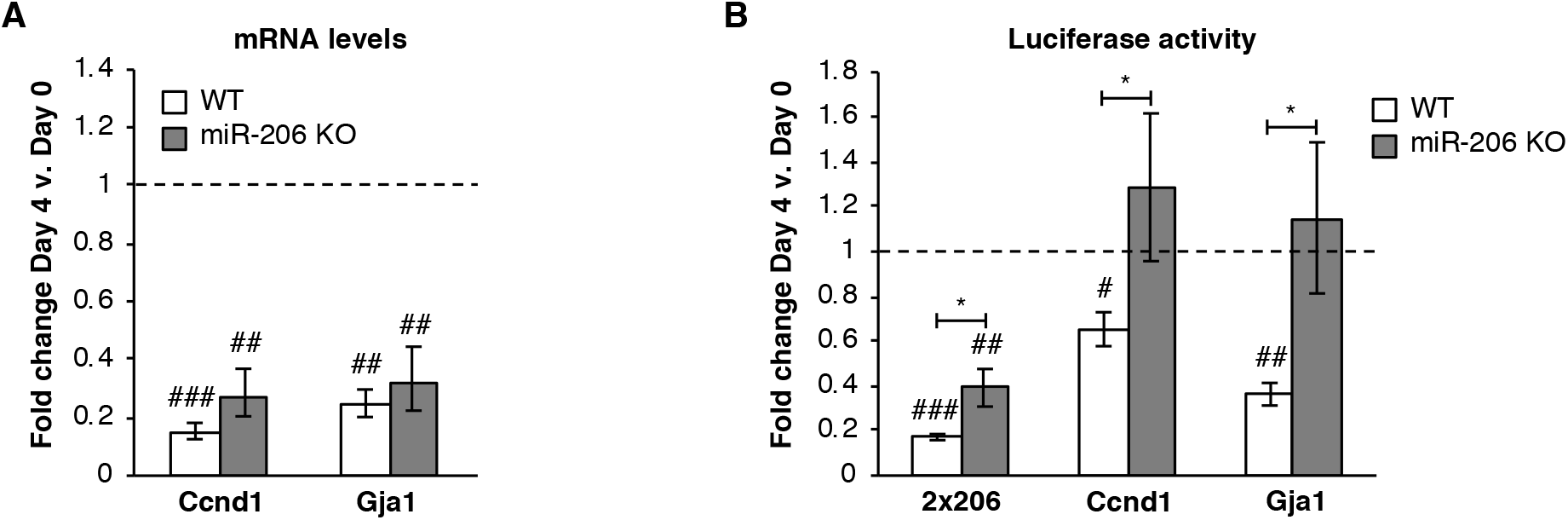
Downregulation of known miR-206 targets is attenuated in miR-206 KO cells compared to WT. (A) Endogenous mRNA levels of Ccnd1 and Gja1 are downregulated in WT cells and miR-206 KO cells after four days in differentiation media. Average fold change in mRNA expression, normalized to Csnk2a2, on Day 4 relative to Day 0 is plotted. The error bars denote standard deviations (n=3). The dashed line at 1.0 represents no change in mRNA levels. ## indicates a p-value < 0.01 and ### indicates p-value < 0.001 as compared to 1.0 (no change), determined from a one sample two-tailed t-test. (B) The 3’UTRs of Ccnd1 and Gja1 downregulate luciferase expression in WT cells but not in miR-206 KO cells upon four days of treatment with differentiation media. Normalized average fold change in luciferase expression after treatment with differentiation media is plotted. The error bars denote standard deviations (n=3). The dashed line at 1.0 represents no change in luciferase expression. # indicates a p-value < 0.05, ## indicates a p-value < 0.01, and ### indicates a p-value < 0.001 as compared to 1.0 using a one sample two-tailed t-test. * indicates a p-value < 0.05, as determined from an unpaired two-tailed t-test comparing the data from the WT and KO cells.

To further investigate the role of miR-206 in regulating Ccnd1 and Gja1, we asked whether their 3’UTRs, each of which contain a miR-206 seed sequence, are sufficient to regulate gene expression in WT or KO cells. We performed reporter assays utilizing plasmids that express transcripts containing the 3’UTR of Gja1 or Ccnd1 fused to the 3’ end of the luciferase RNA (24). Determining if a 3’UTR can control luciferase expression serves as the gold standard in the field for assessing functional miRNA-mRNA binding (29). As a positive control, a third construct (2×206) contained two tandem repeats of the antisense miR-206 sequence fused to the 3’ end of the luciferase RNA (24). These constructs were transfected into WT or miR-206 KO cells, and luciferase expression was measured before and after four days in differentiation media (Figure 2B). In WT cells (white bars), we observed a significant decrease in luciferase expression after four days of differentiation for the 2×206, Ccnd1, and Gja1 constructs, indicating that these 3’UTR sequences were all sufficient to downregulate gene expression during differentiation as miR-206 expression increased. Notably, in the miR-206 KO cells (gray bars), luciferase expression from the Ccnd1 and Gja1 3’UTR constructs no longer decreased when cells were grown in differentiation media. Therefore, regulation of luciferase expression by the Ccnd1 and Gja1 3’UTRs requires miR-206. The 2×206 construct in the miR-206 KO cells showed significantly less repression of luciferase levels compared to the WT cells, revealing a role for miR-206 in downregulating this construct. However, the change in luciferase expression was still significantly less than 1.0; it is likely that miR-1 also targets the 2×206 3’UTR.

### Experimental identification of new mRNA targets of miR-206 in differentiating WT C2C12 cells

To further study miR-206’s role in myogenesis, we were interested in identifying additional miR-206 mRNA targets in differentiating WT C2C12 cells. To do this, we used an experimental affinity purification approach named crosslinking oligo purification (xOP) followed by high-throughput sequencing (xOP-seq; diagrammed in Supplemental Figure 1). In this approach cells were crosslinked with formaldehyde to generate a reversible network of crosslinks, including between miRNA, mRNA, and proteins within miRNA-containing complexes. Cytoplasmic extracts were made, miR-206-containing complexes were captured using a biotinylated DNA oligo antisense to miR-206, and complexes were purified on neutravidin beads. After extensive washing, the crosslinks were reversed, and mRNAs that co-purified with miR-206 in the xOP were identified via Illumina sequencing.

We performed xOP-seq against miR-206 on samples prepared from biological replicates of undifferentiated (Day 0) and Day 4 differentiated WT C2C12 cells. We made 3’end libraries for Illumina sequencing to best capture the 3’UTRs of genes. The sequencing reads were mapped to the mouse genome and analyzed to identify potential miR-206 target mRNAs. First, we bioinformatically identified the 5,512 RNAs that contain at least one miR-206 6mer seed sequence in their 3’UTRs. For this set of RNAs, we quantified the xOP-seq reads that mapped to their 3’UTRs. The mRNAs with at least 2-fold more 3’UTR reads in the Day 4 xOP samples compared to the Day 0 xOP samples, across both biological replicates, were considered potential miR-206 target mRNAs. These criteria yielded 130 putative target mRNAs, which are listed in Supplemental Table 1. We performed gene ontology (GO) enrichment analysis on the list of 130 potential miR-206 targets to identify biological pathways that were significantly enriched. We identified eight pathways that were enriched in the data set (Supplemental Table 2), including intracellular signal transduction, signaling, regulation of response to stimulus, cell communication, and regulation of cellular protein metabolic process. Downregulating components of all these pathways could be important for differentiation.

### MiR-206 regulates expression from several of the newly identified targets

From the list of 130 genes, we chose four for validation experiments: Adam19 (ADAM metallopeptidase domain 19), Bgn (Biglycan), Spg20 (Spastic paraplegia 20), and Tmcc3 (Transmembrane and coiled-coiled domain family 3). These genes all had strong Day 4 xOP-seq peaks in their 3’UTRs, as seen in the sequencing tracks for each gene (Figure 3). Moreover, the peaks are at or near the miR-206 seed sequences. When viewing our xOP-seq data we also found a number of compelling genes that showed 3’UTR peaks near miR-206 seed sequences, which increased in intensity upon differentiation; however, they were not on our candidate list because the Day 4 reads did not exceed the Day 0 reads by at least 2-fold. We chose two additional genes of this type to test in validation experiments: Cbx5 (Chromobox 5) and Smarce1 (SWI/SNF related, matrix associated, actin dependent regulator of chromatin, subfamily E, member 1). In addition to their compelling xOP-seq peaks (see Supplemental Figure 3), downregulation of these genes has the potential to support myogenesis.

**Figure 3.**
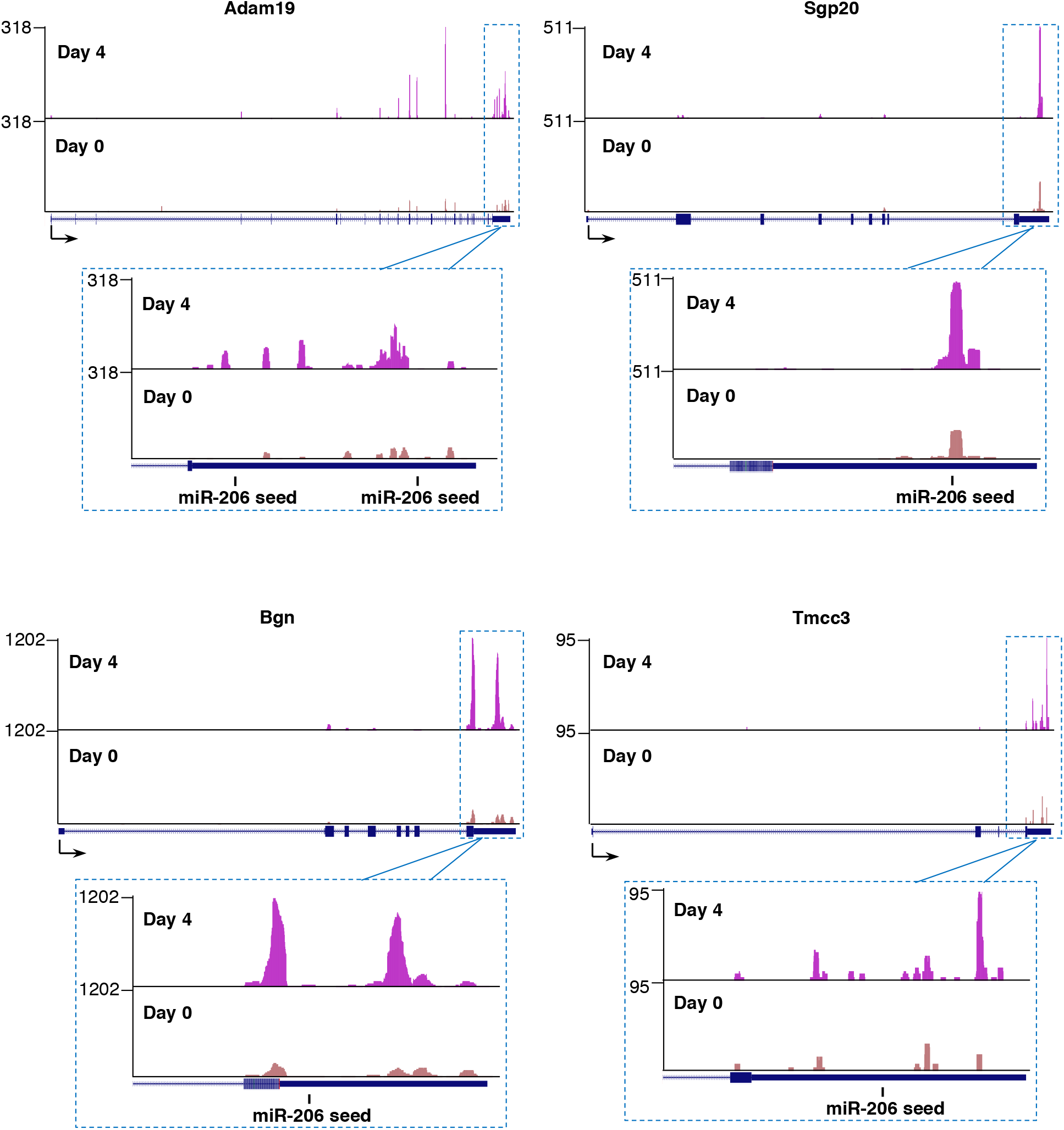
Representative xOP-seq data for Adam19 and Spg20. Mapped reads from sequencing replicate 1 show enrichment in the 3’UTRs in the Day 4 compared to Day 0 data. The locations of miR-206 seed sequences are marked. Sequencing tracks were generated from the UCSC Genome Browser display of bedgraph files.

For the six selected genes, we measured endogenous mRNA levels in both the WT and miR-206 KO cell lines before and four days after growth in differentiation media (Figure 4A). In the WT cells (white bars), four of the six mRNAs significantly decreased upon differentiation: Bgn, Cbx5, Smarce1, and Spg20. In the miR-206 KO cells (gray bars), three of these four mRNAs, Cbx5, Smarce1, and Spg20, did not exhibit significant decreases in their mRNA levels after growth in differentiation media, suggesting that miR-206 has a role in regulating these mRNAs. Adam19 does not exhibit downregulation in either cell line suggesting that if miR-206 regulates expression of Adam19 it must do so at the level of translation and not at level of the transcript. Interestingly, Tmcc3 is upregulated in the WT cells and potentially further unregulated in the miR-206 KO cells upon differentiation. It is possible Tmcc3 is indirectly regulated by miR-206 or that miR-206 is responsible for fine-tuning its expression.

**Figure 4.**
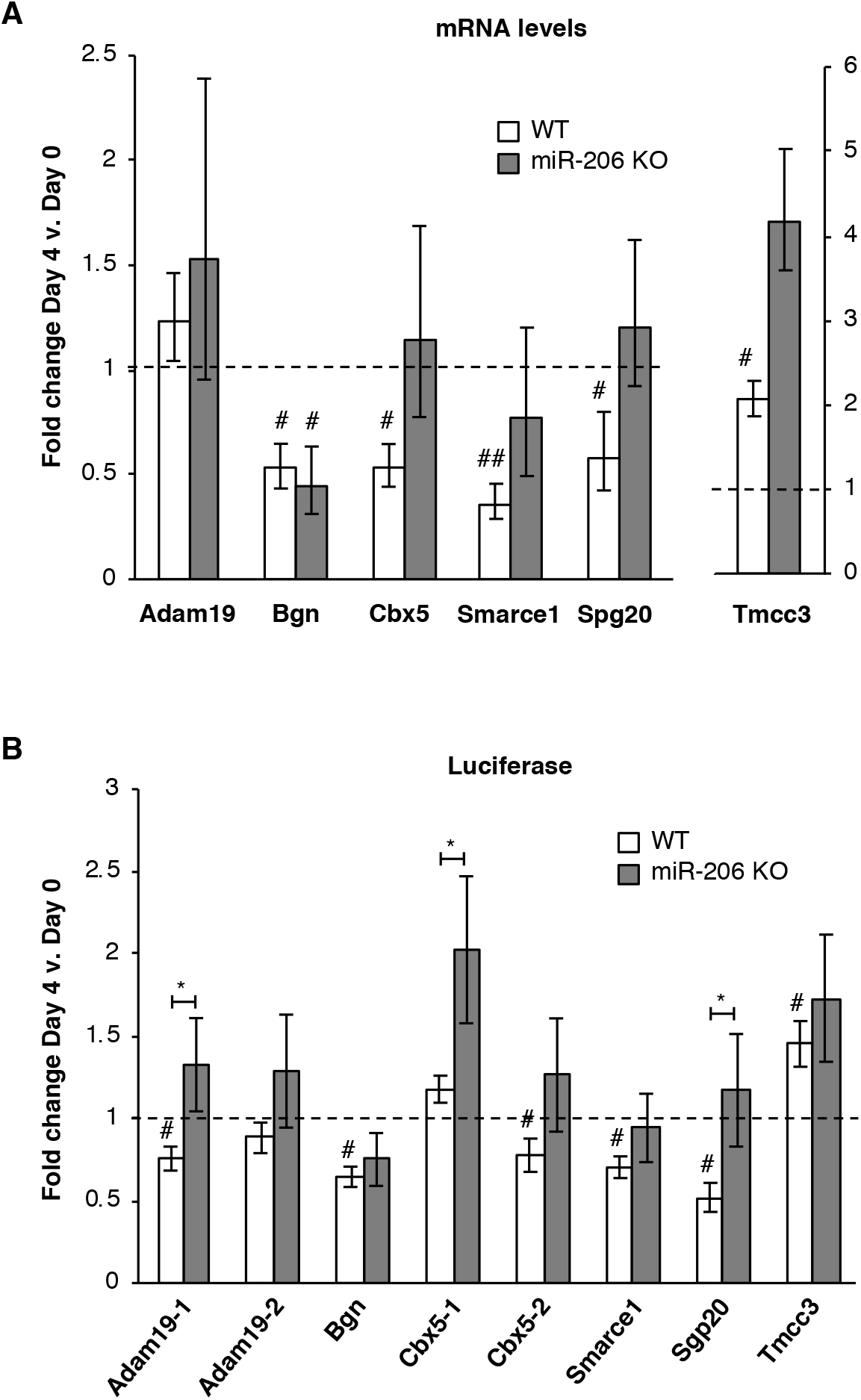
Putative mRNA targets of miR-206 identified by xOP-seq are differentially regulated in miR-206 KO cells compared to WT cells. (A) Endogenous levels of Bgn, Cbx5, Smarce1, and Spg20 mRNAs decrease in WT cells upon differentiation, and show different levels of regulation in miR-206 KO cells after four days in differentiation media. Normalized average fold change in mRNA expression on Day 4 relative to Day 0 is plotted. The error bars denote standard deviations (n=3). The dashed line at 1.0 represents no change in mRNA levels. # indicates a p-value < 0.05 and ## indicates a p-value <0.01 as compared to 1.0 using a one sample two-tailed t-test, except for Spg20 in WT, which has a p-value < 0.068. (B) Regions from the 3’UTRs of five putative miR-206 target mRNAs (Adam19-1, Bgn, Cbx5-1, Smarce1 and Spg20) downregulate luciferase expression in WT cells but not miR-206 KO cells after treatment with differentiation media. Normalized average fold change in luciferase expression after differentiation is plotted; the error bars denote standard deviations (n=3). The dashed line at 1.0 represents no change in luciferase expression. # indicates a p-value < 0.05 as compared to 1.0 using a one sample two-tailed t-test, except for Cbx5-2 in WT, which has a p-value < 0.062. * indicates a p-value < 0.05, as determined from an unpaired two-tailed t-test comparing the data from the WT and KO cells.

For the six candidates, we also constructed reporter plasmids containing each of their 3’UTRs fused to the 3’ end of the luciferase gene. Two constructs were made for Adam19 (Adam19-1, Adam19-2) and Cbx5 (Cbx5-1, Cbx5-2) due to the large size of their 3’UTRs, with each 3’UTR fragment containing a miR-206 seed sequence. We transfected the plasmids into WT or miR-206 KO cells, and measured the fold change in luciferase expression after four days in differentiation media (Figure 4B). In WT cells (white bars), the 3’UTRs from five of the six genes (Adam19-1, Bgn, Cbx5-2, Smarce1, Spg20) caused a statistically significant decrease in luciferase expression upon differentiation, indicative of downregulation as miR-206 expression increased. This is consistent with the mRNA data, and shows that indeed Adam19 is likely regulated by translational inhibition as opposed to mRNA degradation. The 3’UTR fromTmcc3 caused an increase in luciferase expression upon differentiation, much like in the mRNA data. In the miR-206 KO cells (gray bars), all five of the 3’UTR targets that showed decreases in WT cells (Adam19-1, Bgn, Cbx5-2, Smarce1, and Spg20) no longer showed significant decreases in luciferase expression after growth in differentiation media. Moreover, three of the 3’UTR constructs (Adam19-1, Cbx5-1, and Spg20) showed a statistically significant difference between the KO cells and WT cells. Overall, our experiments support the conclusion that Adam19, Bgn, Cbx5, Smarce1, and Spg20 are bona fide miR-206 targets, highlighting the utility of the xOP-seq method for experimentally identifying mRNA targets of miRNAs.

## Discussion

Here we used CRISPR technology to knock out the gene encoding miR-206 in C2C12 cells. We confirmed that the KO cells did not express miR-206, but still expressed normal levels of the related microRNA miR-1 after treatment with differentiation media. The miR-206 KO cell line was delayed and deficient in differentiation, and did not form myotubes, although 2 differentiation markers were expressed at normal levels in the KO cells. An experimental method (xOP-seq) was used to identify over a hundred potential new mRNA targets of miR-206 in C2C12 cells. The majority of identified mRNAs tested experimentally were found to be direct targets of miR-206.

We found that knocking out miR-206 in C2C12 myoblasts dramatically decreased their differentiation capacity and ability to form myotubes. This is consistent with previous studies that introduced antagomiRs (i.e. oligos antisense to miRNAs) into cells to transiently perturb regulation by miR-206. For example, antagomiRs against miR-206, miR-1, and miR-133 in combination caused a shortening of myotubes upon differentiation of C2C12 cells (12). Similarly, C2C12 cells transfected with a miR-206 antagomiR, which targeted both miR-206 and miR-1, had a reduced fusion index and total myotube area (18). While our data are consistent with these findings, our approach provides significant new insight. Importantly, by using CRISPR/Cas9 technology to provide a genetic perturbation, we ensured only miR-206 was knocked out and remained so throughout days of differentiation. Uniquely, this enabled us to study the specific role of miR-206, without perturbing miR-1, in a useful cell culture model of myogenesis. Our phenotypic analysis of the KO cell line demonstrates that miR-206 alone has important roles in skeletal muscle development. Specifically, it impacts myoblast fusion and delays myotube formation and elongation. The molecular mechanisms by which myotube formation is disrupted in the miR-206 KO cells is not known. However, we show that myosin and myogenin protein levels were not affected in the miR-206 KO compared to WT; therefore, the delayed differentiation phenotypes in the KO cells are not attributable to changing levels of these important differentiation markers. Prior studies showed that miR-206/1 antagomiRs decreased myosin heavy chain levels upon differentiation (12, 18); however, this could be due to miR-1 perturbation with the use of antagomiRs.

Our finding that miR-206 itself is critical for myogenesis in the C2C12 model system is consistent with the handful of *in vivo* miR-206 KO mouse studies showing miR-206 is important in muscle biology. Although knocking out miR-206 in mice does not result in a lethal phenotype, these mice experience a number of abnormalities including reduced satellite cell differentiation, delayed muscle regeneration, as well changes to their skeletal and cardiac slow muscle programming (19, 30). Disease state mouse models of DMD and ALS lacking miR-206 show exacerbated disease progression (19, 20). Profound effects on neuromuscular junction reinnervation after injury were observed in the ALS model (20), and impaired muscle regeneration, integrity, and function were observed in the DMD model (19). Perhaps we can better understand the contribution of miR-206 to these disease states by better understanding how it functions in normal muscle cells. The miR-206 KO cell culture model we generated provides an experimentally tractable system for future mechanistic studies to investigate the molecular basis for the contributions miR-206 makes to muscle differentiation.

xOP-seq identified a set of 130 potential miR-206 targets in differentiating C2C12 cells. Some, but not all, known miR-206 targets were part of this set, including HDAC4 (31), Igfbp5 (19), and Map1a (32). The criterion we used to obtain our list of 130 putative miR-206 targets required a 2-fold or greater enrichment of xOP-seq reads in the differentiated sample. Known targets like Gja1 and Ccnd1 did not meet this criteria and it is possible there are other potential bona fide miR-206 targets that did not meet this stringent cut-off as well. Indeed, we experimentally tested two mRNAs recovered from xOP-seq – Cbx5 and Smarce1 – for which this was true, and the experimental validation data support both being miR-206 targets. In the future, xOP-seq could include a means to quantify absolute transcript levels in each sample to identify putative target enrichment, rather than comparing relative sequencing counts.

We selected six mRNAs to test experimentally as putative miR-206 targets. Two types of experiments were performed. First, we assessed endogenous mRNA levels before and after treating WT and miR-206 KO cells with differentiation media (Figure 4A). Four of the mRNAs (Bgn, Cbx5, Smarce1, and Spg20) showed downregulation after four days of differentiation in WT cells, and three of these were no longer downregulated in the KO cells. Second, we tested whether the 3’UTRs from the six mRNAs could regulate luciferase expression in WT and KO cells treated with differentiation media. For Adam19, Cbx5, Smarce1, Spg20, and to a lesser extent Bgn, we observed downregulation of luciferase levels in WT cells, coupled to an attenuation downregulation in the miR-206 KO cell line (Figure 4B). For some target mRNAs, our data showed discrepancies between the level of endogenous mRNA downregulation and the level of downregulation in the 3’UTR luciferase assay. This was particularly true for the known miR-206 targets Ccnd1 and Gja1, as well as for the newly identified miR-206 target Bgn. In the 3’UTR luciferase assays these targets showed downregulation in the WT cells that was alleviated in the miR-206 KO. However, these targets showed significant endogenous mRNA downregulation in both the WT and the miR-206 KO cell line. This suggests these targets experience an additional layer of regulation outside their 3’UTRs upon differentiation that is unrelated to miR-206, perhaps due to another miRNA targeting the mRNAs in their 5’UTRs or coding regions (33–35). Together our data support the conclusion that Adam19, Bgn, Cbx5, Smarce1, and Spg20 are novel miR-206 targets

These newly identified mRNA targets have interesting connections to skeletal muscle differentiation. Cbx5 and Smarce1 are both chromatin regulators and terminal differentiation of skeletal muscle involves chromatin reorganization (36). Cbx5 (Hp1α) can interact with and inhibit the activity of MyoD, a necessary transcription factor in myogenic commitment (37). Cbx5 is also known to associate with HDAC4 (38), a known target of miR-206 that represses another pro-differentiation skeletal muscle transpiration factor, MEF2 (39). Smarce1 (BAF57) cooperates with zinc-finger containing factor TSHZ3 to silence MyoD-dependent myogenin expression (40). Therefore, downregulation of Cbx5 and Smarce1 by miR-206 would support the differentiation programing by allowing important myogenic transcription factors to be active. Bgn expression has previously been shown to decrease during skeletal muscle differentiation, consistent with our results (41). In the future, it will be interesting to further study the myogenic roles of these targets. While we only chose a handful of putative miR-206 targets to validate, further studies of other xOP-seq identified genes could reveal additional direct miR-206 targets.

## Methods

### Cell culture

C2C12 myoblast cells were cultured in 5% CO2 at 37°C and maintained at ~70% confluency in DMEM supplemented with 20% FBS, 1% penicillin-streptomycin, 1% sodium pyruvate, and 1% L-glutamate (growth media). Differentiation was induced by culturing cells in differentiation media (DMEM supplemented with 5% horse serum, 1% penicillin-streptomycin, 1% sodium pyruvate, and 1% L-glutamate).

### Generating a miR-206 knock-out in C2C12 cells using CRISPR/Cas9

An Alt-R CRISPR-Cas9 crRNA was designed to cut in the seed of the mature miR-206 sequence (5’-TCTCAGCACTATGGAATGTA-3’; IDT). We used an Alt-R CRISPR-Cas9 Control Kit Mouse (IDT), which included Alt-R CRISPR tracrRNA, a positive control crRNA to the HPRT gene, and a negative control crRNA. All crRNAs were annealed to tracrRNA to create a crRNA:tracrRNA duplex. C2C12 WT myoblasts were seeded at a density of 25,000 cells/well in 6-well plates (Corning Costar). 24 hours after seeding, 2.5 μg of pSpCas9(BB)-2A-GFP (PX458, Addgene) plasmid was transfected into cells using Lipofectamine 3000 (ThermoFischer Scientific). 48 hours after seeding, 3 μM crRNA:tracrRNA duplex (miR-206, HPRT, or negative control) was transfected into cells using INTERFERin (Polyplus transfection). 72 hours after seeding, GFP expressing cells were sorted by FACS and checked for overall Cas9 targeting efficiency using T7E1 assays on isolated genomic DNA. After observing specific cutting by Cas9, the transfection and sorting were repeated to isolate single cells expressing Cas9-GFP, which were placed into 96-well plates and passaged. A control clonal cell line transfected with the negative control crRNA:tracrRNA duplex was created in parallel. Genomic DNA was extracted from single clonal populations of cells, and the miR-206 target region was amplified by PCR, cloned into pUC18, and sequenced to identify INDELs.

### Jenner-Giemsa staining

C2C12 WT and miR-206 KO cells were seeded at a density of 50,000 cells/well in 6-well plates (Corning Costar). Cells were cultured in growth media for 48 hours before switching to differentiation media. Time points were as follows: 24 hours in growth media (Day −1); 48 hours in growth media (Day 0); 24, 48, 96, and 144 hours in differentiation media (Days 1, 2, 4, 6, respectively). Differentiation media was changed every 48 hours. Cells were stained in biological triplicate as described in (28). Briefly, cells were fixed with 100% methanol, stained with a 1:3 dilution of Jenner staining solution (Electron Microscopy Sciences), then stained with a 1:20 dilution of Giemsa staining solution (Electron Microscopy Sciences). Stained cells were imaged using an Evos FL Imaging System (Invitrogen) and a Nikon SMZ25 Stereomicroscope. For the data in Figure 1C, images were quantified as described in (28).

### RT-qPCR

C2C12 WT and miR-206 KO myoblasts were seeded at a density of 50,000 cells/well in 6-well plates (Corning Costar), cultured in growth media for 48 hours (Day 0), then switched to differentiation media for 96 hours (Day 4). Differentiation media was changed every 48 hours. Total RNA was extracted from Day 0 and Day 4 cells in biological triplicate using Ribozol (Amresco), then DNaseI treated with TURBO DNase (Invitrogen).

For miRNA analysis, the TaqMan MicroRNA Reverse Transcription Kit (Applied Biosystems) was used along with the following specific RT stem-loop primers: hsa-miR-206, Assay ID: 000510; hsa-miR-1, Assay ID: 002222; sno202, Assay ID: 001232. The expression level of each miRNA was normalized to sno202 levels, and the fold-changes in expression between WT and miR-206 KO cells were calculated using the ΔΔCT method.

For mRNA analysis, RNA was reverse transcribed using 6 μM random hexamers, 500 μM dNTPs, 10 μM DTT, 1 U RNase Inhibitor, Murine (NEB), and 5 U M-MuLV RT (NEB) in RT Buffer (25 mM KCl, 50 mM Tris-HCl, pH 7.5, 3.5 mM MgCl_2_, 0.1 mg/mL BSA). Reactions were incubated at 25°C for 5 min, 42°C for 60 minutes, 65°C for 20 minutes, then held at 4°C. PCR was performed in technical duplicates using a StepOnePlus Real-Time PCR system (Applied Biosystems) with SYBR Green detection. The expression level of each mRNA was normalized to Csnk2a2 and the fold-changes between WT and KO cells were calculated using the ΔΔCT method. PCR primer sequences are in Supplementary Table 3.

### 3’UTR luciferase reporter assays

Luciferase reporter vectors were made by PCR amplifying the entire 3’UTR (or in the case of Adam19 and Cbx5, two 3’UTR segments) from mouse genomic DNA using primers with XhoI and NotI restriction sites. Using these cut sites, the PCR fragments were cloned into the psiCHECK2 vector (Promega) downstream of *Renilla* luciferase. This vector also contains firefly luciferase as an internal control. psiCHECK-EMPTY, psiCHECK-2×206 and psiCHECK2-*Ccnd1* 3’UTR vector constructs are described elsewhere (24).

psiCHECK2 vector constructs were transfected into WT and miR-206 knock-out C2C12 myoblasts 24 hours after seeding at a density of 50,000 cells/well in 6-well plates with growth media. Cells were switched to differentiation media 48 hours after seeding. Cells were harvested after 48 hours in growth media (Day 0), and 96 hours in differentiation media (Day 4). Differentiation media was changed every 48 hours. Luminescence of *Renilla* and firefly luciferase was measured using the Dual-Luciferase Reporter Assay System (Promega, E1910) using a Synergy H1 Hybrid Multi-Mode Microplate Reader (BioTek). *Renilla* luciferase activity was normalized by the control firefly luciferase for each construct on Day 0 and Day 4. This *Renilla/firefly* ratio for each 3’ UTR construct was then normalized to the *Renilla/firefly* ratio for the psiCHECK2-EMPTY vector (a negative control construct containing no 3’UTR) on Day 0 and Day 4. The fold change in normalized luciferase activity between Day 4 and Day 0 was then calculated for each 3’UTR construct.

### Crosslinking-oligo purification (xOP)

C2C12 myoblasts were seeded in 15 cm plates (Corning) at a density of 75,000 cells. After 48 hours in growth media, cells were either crosslinked and harvested (undifferentiated/Day 0) or switched to differentiation media for 96 hours before crosslinking and harvesting (Day 4 differentiated). For each condition, cells from 27 plates in duplicate were crosslinked with 1% formaldehyde, shaking for 10 minutes at room temperature, before quenching with glycine (0.125 M, shaking for 10 minutes at room temperature). Cells were washed two times with ice-cold 1X PBS, scraped in 1X ice-cold PBS with 1 mM PMSF, pelleted and stored at −80°C until lysis.

To generate cytoplasmic extracts, cells were resuspended in 5 volumes of lysis buffer (20 mM HEPES pH 7.9, 1.5 mM MgCl_2_, 10 mM KCl, 1 mM DTT, 1 mM PMSF, 1X Protease Inhibitor) and incubated on ice for 20 minutes. Using an ice-cold dounce homogenizer, cells were dounced 30 times with a type B pestle. Samples were pelleted at maximum speed, the supernatant collected, and 300 mM KCl, 100 mM EDTA, and 0.1% NP-40 were added. The extracts were pre-cleared with NeutrAvidin agarose resin (Pierce) prepared by washing three times with xOP Buffer (20 mM HEPES pH 7.9, 2 mM EDTA, 300 mM KCl, 1 mM DTT, 1 mM PMSF, 0.2% NP-40). 20 μL packed beads/1 mL cytoplasmic extract were added and nutated for 1 hour at 4°C. The beads were spun down at 2000 rpm for 2 minutes at 4°C and the supernatant was collected. Ten percent of the pre-cleared cytoplasmic extracts were saved as input samples.

For the xOP pull down, 600 pmol of a miR-206 biotinylated DNA antisense oligo (5’ biotinTEG-CCACACACTTCCTTACATTCCA-3’) (IDT) was added to pre-cleared cytoplasmic extracts. Samples were heated to 95°C for 5 minutes, slow cooled to room temperature, and nutated at 4°C for 20-30 minutes. High Capacity NeutrAvidin agarose resin (Pierce) was prepared by washing three times with xOP Buffer. The prepared beads were added (20 μL packed beads/1 mL cytoplasmic extract) and nutated for 1 hour at 4°C. The beads were washed with 15 bead volumes as follows: twice with xOP Buffer, once with High Salt Buffer (20 mM HEPES pH 7.9, 2 mM EDTA, 1 M KCl, 1 mM DTT, 1 mM PMSF, 0.2% NP-40), once with LiCl Buffer (20 mM HEPES pH 7.9, 1 mM EDTA, 250 mM LiCl, 1 mM DTT, 1% deoxycholate, 1% NP-40), twice with Pre-Elution Buffer (50 mM HEPES pH 7.9, 5 mM EDTA, 200 mM KCl, 1 mM DTT, 0.5% NP-40), and once with 2 bead volumes of 1X RNase H Buffer (NEB; 50mM Tris-HCl, 3 mM MgCl_2_, 10 mM DTT, pH 8.3). 100 μL of 1X RNase H Buffer was added to transfer the beads to a new tube before spinning down and removing the final wash. To elute RNA, 50 μL of 1X RNase H Buffer and 5μL of RNase H (NEB) were added to the beads and incubated at 37°C for 30 minutes while nutating. The supernatant was collected as eluate. 62.5μL of extra buffer solution (50 mM HEPES pH 7.9, 5 mM EDTA, 1 mM DTT) was added to the beads and incubated at 37°C for 5 minutes for a final elution. The supernatant was collected and added to the previously collected eluate. 331.5 μL of extra buffer solution (50 mM HEPES pH 7.9, 5 mM EDTA, 1 mM DTT) with 300 mM KCl was added to bring the final volume up to 500 μL. Proteinase K (100 μg/mL) was added to the eluate and input samples and heated at 65°C for 1 hour. RNA was extracted using the miRNeasy Mini Kit (Qiagen) and DNaseI treated with TURBO DNase (Invitrogen).

### Library preparation and high throughput RNA-sequencing

Libraries for Illumina Sequencing were prepared as follows. RNA eluates were PolyC tailed in 20 μL reactions at 37°C for 15 minutes using 1 mM CTP, 0.1 U PolyU polymerase (NEB), 0.25 U *E. coli* PolyA polymerase (NEB), and 1 U RiboLock RNase Inhibitor (ThermoFisher Scientific) in 1X *E. coli* Poly(A) Polymerase Reaction Buffer (NEB) and 1X NEBuffer 2. RNA was isolated using Ribozol (Amresco), then annealed to Illumina compatible primer for reverse transcription (1 μM; 5’-AGTTCAGACGTGTGCTCTTCCGATCTGGGGGGGGGGGHN-3’) in 7 mM Tris-HCl, pH 7.5, 35 mM KCl and 0.75 mM EDTA by heating to 95°C for 5 min, then slow cooling to 48°C and holding for 30 minutes. Samples were reverse transcribed using 500 μM dNTPs, 10 μM DTT, 1 U RiboLock RNase Inhibitor (ThermoFisher Scientific), and 1 U MultiScribe RT (Invitrogen) in RT Buffer (25 mM KCl, 50 mM Tris-HCl, pH 7.5, 3.5 mM MgCl_2_, 0.1 mg/mL BSA) and incubating at 42°C for 60 minutes, 85°C for 5 minutes, then holding at 4°C. Samples were RNaseH treated and cleaned up using the E.Z.N.A Cycle-Pure Kit (Omega Bio-Tek). Second strand cDNA was prepared by incubating 0.5 μM dNTPs, 1 μM Illumina-compatible random hexamer primer (5’-TCCCTACACGACGCTCTTCCGATCTNNNNNN-3’), and 0.15 U Klenow Fragment (3’→5’ exo-) (NEB) in 1X NEBuffer 2 at 37°C for 30 minutes (cite). Double stranded DNA was purified using the E.Z.N.A Cycle-Pure Kit (Omega Bio-Tek) then PCR amplified for 13 cycles using 200 μM dNTPs, 240 μM TruSeq Indexed Adapters (AD001; AD003; AD008; AD009), 240 μM TruSeq Universal Adapter (5’-AATGATACGGCGACCACCGAGATCTACACTCTTTCCCTACACGACGCTCTTCCGATCT-3’) and 0.025 U *Taq* DNA Polymerase (NEB) in 1X Standard *Taq* Reaction Buffer (NEB). The libraries were gel-purified using the E.Z.N.A Gel Extraction Kit (Omega Bio-Tek). Sequencing was performed using a HiSeq 4000, obtaining 1×50 bp reads with a 15% PhiX spike-in.

### Computational analysis of sequencing reads

Reads were mapped to the mouse genome (mm10 assembly) using Bowtie2 (version 2.3.2) in the -D 5 -R 1 -N 0 -L 25 -i S,1,2.00 alignment mode plus trimming 10 bps off the 3’ end where read quality decreased. Meta-analyses and per-gene quantification of mapped reads were performed using the suites of tools available in HOMER (Hypergeometric Optimization of Motif EnRichment) and in Bedtools (v2.26.0). Tag directories in HOMER were created from the sam files generated by Bowtie2. To allow visualization of the mapped reads in the UCSC Genome Browser, bedGraph files were generated using the makeUCSCfile command in HOMER. The bedtools intersect command in Bedtools was used to identify RefSeq genes that contained a miR-206 6mer seed (5’-CCTTAC-3’) in their 3’UTR. The tag directories created in HOMER were converted to bed files using the tagDir2bed command in HOMER. The bedtools coverage command in Bedtools was used to count RNA sequence tags across the 3’UTRs of genes that contained a miR-206 6mer seed. Reads were normalized to a constant depth of sequencing for each data set and divided by the length of the 3’UTR of each gene to obtain reads per kilobase (RPK). Genes with RPK values for Day 4 xOP that were at least 2-fold above Day 0 xOP for each replicate defined the set of putative miR-206 targets. Gene ontology analyses were performed using the Functional Annotation tool in DAVID Bioinformatics Resources 6.8, with the gene ontology category GOTERM_BP_ALL. Enriched terms had Benjamini-Hochberg corrected p-values of ≤ 0.1.

## Acknowledgements

Wild-type C2C12 cells, psiCHECK2-EMPTY, psiCHECK2-2×206, and psiCHECK-*Ccnd1* 3’UTR plasmid were generous gifts from Dr. Kristen K. Bjorkman and the Leinwand Lab at CU Boulder. The authors would also like to thank Dr. Bjorkman for all her invaluable help, advice, and discussion throughout this work. This work was supported by grants R21 AR06782 and R01 GM068414 from the National Institutes of Health. G.M.S was partially supported by training grant T32 GM08759 from the National Institutes of Health.

## References

1. Bartel, D. P. (2004) MicroRNAs: Genomics, Biogenesis, Mechanism, and Function. Cell. 116, 281–297

2. Bartel, D. P. (2009) MicroRNA Target Recognition and Regulatory Functions. Cell. 136, 215–233

3. Kloosterman, W. P., and Plasterk, R. H. A. (2006) The Diverse Functions of MicroRNAs in Animal Development and Disease. Dev. Cell. 11, 441–450

4. Almeida, M. I., Reis, R. M., and Calin, G. A. (2011) MicroRNA history: Discovery, recent applications, and next frontiers. Mutat. Res. Mol. Mech. Mutagen. 717, 1–8

5. Chen, J.-F., Callis, T. E., and Wang, D.-Z. (2009) microRNAs and muscle disorders. J. Cell Sci. 122, 13–20

6. MacFarlane, L.-A., and Murphy, P. R. (2010) MicroRNA: Biogenesis, Function and Role in Cancer. Curr. Genomics. 11, 537–561

7. O’Rourke, J. R., Georges, S. A., Seay, H. R., Tapscott, S. J., McManus, M. T., Goldhamer, D. J., Swanson, M. S., and Harfe, B. D. (2007) Essential role for Dicer during skeletal muscle development. Dev. Biol. 311, 359–368

8. Bentzinger, C. F., Wang, Y. X., and Rudnicki, M. A. (2012) Building Muscle: Molecular Regulation of Myogenesis. Cold Spring Harb. Perspect. Biol. 10.1101/cshperspect.a008342

9. Horak, M., Novak, J., and Bienertova-Vasku, J. (2016) Muscle-specific microRNAs in skeletal muscle development. Dev. Biol. 410, 1–13

10. Goljanek-Whysall, K., Sweetman, D., and Münsterberg, A. E. (2012) microRNAs in skeletal muscle differentiation and disease. Clin. Sci. 123, 611–625

11. Chen, J.-F., Mandel, E. M., Thomson, J. M., Wu, Q., Callis, T. E., Hammond, S. M., Conlon, F. L., and Wang, D.-Z. (2006) The role of microRNA-1 and microRNA-133 in skeletal muscle proliferation and differentiation. Nat. Genet. 38, 228–233

12. Kim, H. K., Lee, Y. S., Sivaprasad, U., Malhotra, A., and Dutta, A. (2006) Muscle-specific microRNA miR-206 promotes muscle differentiation. J. Cell Biol. 174, 677–687

13. Rao, P. K., Kumar, R. M., Farkhondeh, M., Baskerville, S., and Lodish, H. F. (2006) Myogenic factors that regulate expression of muscle-specific microRNAs. Proc. Natl. Acad. Sci. U. S. A. 103, 8721–8726

14. Koutsoulidou, A., Mastroyiannopoulos, N. P., Furling, D., Uney, J. B., and Phylactou, L. A. (2011) Expression of miR-1, miR-133a, miR-133b and miR-206 increases during development of human skeletal muscle. BMC Dev. Biol. 11, 34

15. Sweetman, D., Goljanek, K., Rathjen, T., Oustanina, S., Braun, T., Dalmay, T., and Münsterberg, A. (2008) Specific requirements of MRFs for the expression of muscle specific microRNAs, miR-1, miR-206 and miR-133. Dev. Biol. 321, 491–499

16. Sweetman, D., Rathjen, T., Jefferson, M., Wheeler, G., Smith, T. G., Wheeler, G. N., Münsterberg, A., and Dalmay, T. (2006) FGF-4 signaling is involved in mir-206 expression in developing somites of chicken embryos. Dev. Dyn. 235, 2185–2191

17. Zhao, Y., Samal, E., and Srivastava, D. (2005) Serum response factor regulates a muscle-specific microRNA that targets Hand2 during cardiogenesis. Nature. 436, 214–220

18. Goljanek-Whysall, K., Pais, H., Rathjen, T., Sweetman, D., Dalmay, T., and Münsterberg, A. (2012) Regulation of multiple target genes by miR-1 and miR-206 is pivotal for C2C12 myoblast differentiation. J. Cell Sci. 125, 3590–3600

19. Liu, N., Williams, A. H., Maxeiner, J. M., Bezprozvannaya, S., Shelton, J. M., Richardson, J. A., Bassel-Duby, R., and Olson, E. N. (2012) microRNA-206 promotes skeletal muscle regeneration and delays progression of Duchenne muscular dystrophy in mice. J. Clin. Invest. 122, 2054–2065

20. Williams, A. H., Valdez, G., Moresi, V., Qi, X., McAnally, J., Elliott, J. L., Bassel-Duby, R., Sanes, J. R., and Olson, E. N. (2009) MicroRNA-206 Delays ALS Progression and Promotes Regeneration of Neuromuscular Synapses in Mice. Science. 326, 1549–1554

21. Alteri, A., De Vito, F., Messina, G., Pompili, M., Calconi, A., Visca, P., Mottolese, M., Presutti, C., and Grossi, M. (2013) Cyclin D1 is a major target of miR-206 in cell differentiation and transformation. Cell Cycle. 12, 3781–3790

22. Anderson, C., Catoe, H., and Werner, R. (2006) MIR-206 regulates connexin43 expression during skeletal muscle development. Nucleic Acids Res. 34, 5863–5871

23. Rosenberg, M. I., Georges, S. A., Asawachaicharn, A., Analau, E., and Tapscott, S. J. (2006) MyoD inhibits Fstl1 and Utrn expression by inducing transcription of miR-206. J. Cell Biol. 175, 77–85

24. Bjorkman, K. K., Buvoli, M., Pugach, E. K., Polmear, M. M., and Leinwand, L. A. (2019) miR-1/206 downregulates splicing factor Srsf9 to promote C2C12 differentiation. Skelet. Muscle. 9, 31

25. Ma, G., Wang, Y., Li, Y., Cui, L., Zhao, Y., Zhao, B., and Li, K. (2015) MiR-206, a Key Modulator of Skeletal Muscle Development and Disease. Int. J. Biol. Sci. 11, 345–352

26. Chen, J.-F., Tao, Y., Li, J., Deng, Z., Yan, Z., Xiao, X., and Wang, D.-Z. (2010) microRNA-1 and microRNA-206 regulate skeletal muscle satellite cell proliferation and differentiation by repressing Pax7. J. Cell Biol. 190, 867–879

27. Dey, B. K., Gagan, J., and Dutta, A. (2011) miR-206 and -486 Induce Myoblast Differentiation by Downregulating Pax7. Mol. Cell. Biol. 31, 203–214

28. Veliça, P., and Bunce, C. M. (2011) A quick, simple and unbiased method to quantify C2C12 myogenic differentiation. Muscle Nerve. 44, 366–370

29. Nicolas, F. E. (2011) Experimental Validation of MicroRNA Targets Using a Luciferase Reporter System. in MicroRNAs in Development: Methods and Protocols (Dalmay, T. ed), pp. 139–152, Methods in Molecular Biology, Humana Press, Totowa, NJ, 10.1007/978-1-61779-083-6_11

30. Bjorkman, K. K., Guess, M. G., Harrison, B. C., Polmear, M. M., Peter, A. K., and Leinwand, L. A. (2019) miR-206 Enforces a Slow Muscle Phenotype. bioRxiv. 10.1101/756981

31. Winbanks, C. E., Wang, B., Beyer, C., Koh, P., White, L., Kantharidis, P., and Gregorevic, P. (2011) TGF-β Regulates miR-206 and miR-29 to Control Myogenic Differentiation through Regulation of HDAC4. J. Biol. Chem. 286, 13805–13814

32. Tang, Z., Yang, Y., Wang, Z., Zhao, S., Mu, Y., and Li, K. (2015) Integrated analysis of miRNA and mRNA paired expression profiling of prenatal skeletal muscle development in three genotype pigs. Sci. Rep. 5, 15544

33. Lytle, J. R., Yario, T. A., and Steitz, J. A. (2007) Target mRNAs are repressed as efficiently by microRNA-binding sites in the 5’ UTR as in the 3’ UTR. Proc. Natl. Acad. Sci. 104, 9667–9672

34. Tay, Y., Zhang, J., Thomson, A. M., Lim, B., and Rigoutsos, I. (2008) MicroRNAs to Nanog, Oct4 and Sox2 coding regions modulate embryonic stem cell differentiation. Nature. 455, 1124–1128

35. Forman, J. J., Legesse-Miller, A., and Coller, H. A. (2008) A search for conserved sequences in coding regions reveals that the let-7 microRNA targets Dicer within its coding sequence. Proc. Natl. Acad. Sci. U. S. A. 105, 14879–14884

36. Harada, A., Ohkawa, Y., and Imbalzano, A. N. (2017) Temporal Regulation of Chromatin During Myoblast Differentiation. Semin. Cell Dev. Biol. 72, 77–86

37. Yahi, H., Fritsch, L., Philipot, O., Guasconi, V., Souidi, M., Robin, P., Polesskaya, A., Losson, R., Harel-Bellan, A., and Ait-Si-Ali, S. (2008) Differential Cooperation between Heterochromatin Protein HP1 Isoforms and MyoD in Myoblasts. J. Biol. Chem. 283, 23692–23700

38. Zhang, C. L., McKinsey, T. A., and Olson, E. N. (2002) Association of Class II Histone Deacetylases with Heterochromatin Protein 1: Potential Role for Histone Methylation in Control of Muscle Differentiation. Mol. Cell. Biol. 22, 7302–7312

39. Lu, J., McKinsey, T. A., Zhang, C.-L., and Olson, E. N. (2000) Regulation of Skeletal Myogenesis by Association of the MEF2 Transcription Factor with Class II Histone Deacetylases. Mol. Cell. 6, 233–244

40. Faralli, H., Martin, E., Coré, N., Liu, Q.-C., Filippi, P., Dilworth, F. J., Caubit, X., and Fasano, L. (2011) Teashirt-3, a Novel Regulator of Muscle Differentiation, Associates with BRG1-associated Factor 57 (BAF57) to Inhibit Myogenin Gene Expression. J. Biol. Chem. 286, 23498–23510

41. Casar, J. C., McKechnie, B. A., Fallon, J. R., Young, M. F., and Brandan, E. (2004) Transient up-regulation of biglycan during skeletal muscle regeneration: delayed fiber growth along with decorin increase in biglycan-deficient mice. Dev. Biol. 268, 358–371

