## Supplementary Materials for "miR-206 knockout shows it is critical for myogenesis and directly regulates newly identified target mRNAs"

Supplemental Figures 1-2
Supplemental Tables 1-3

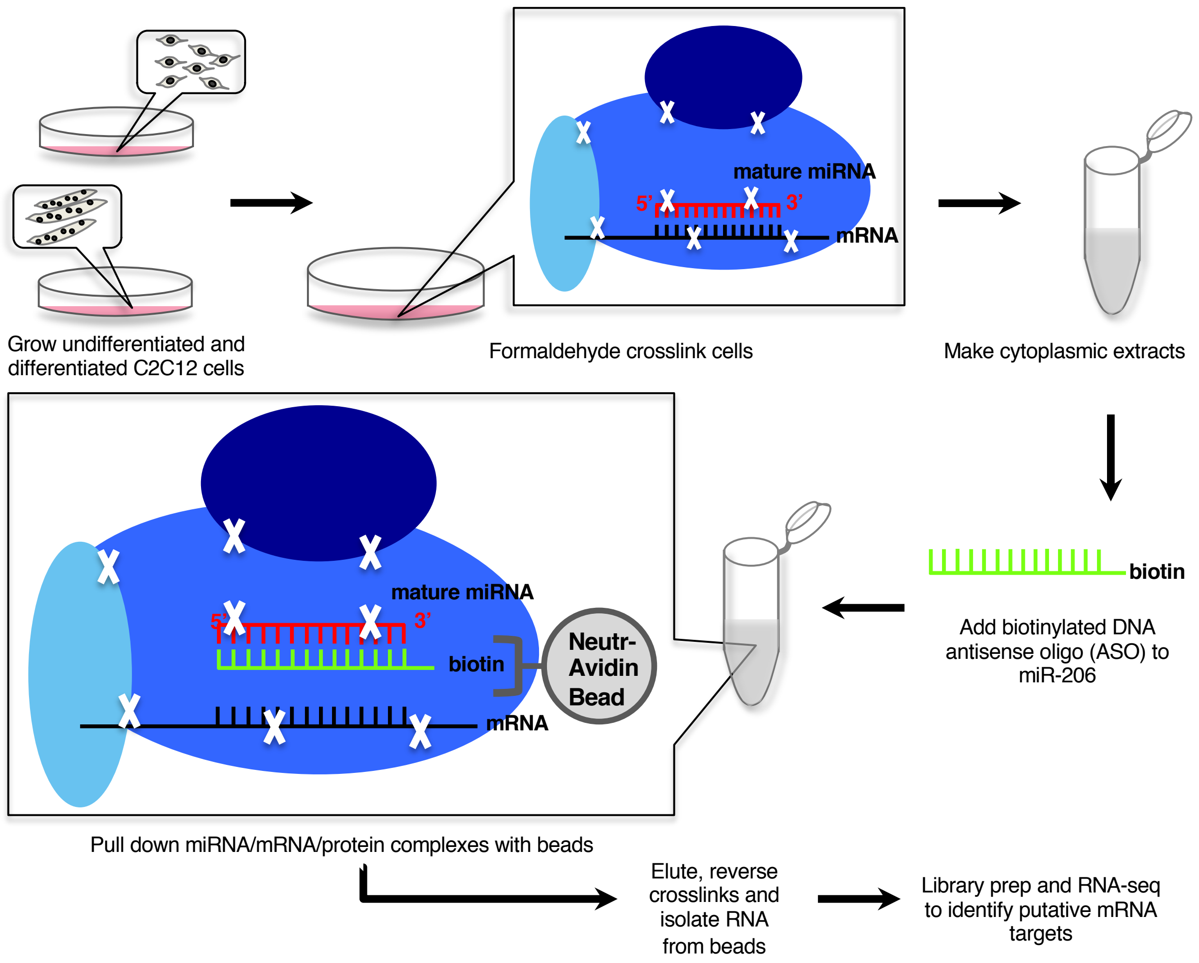


**Supplemental Figure 1. Schematic of xOP-seq method workflow.** See text for a complete description. The white X's represent crosslinks.

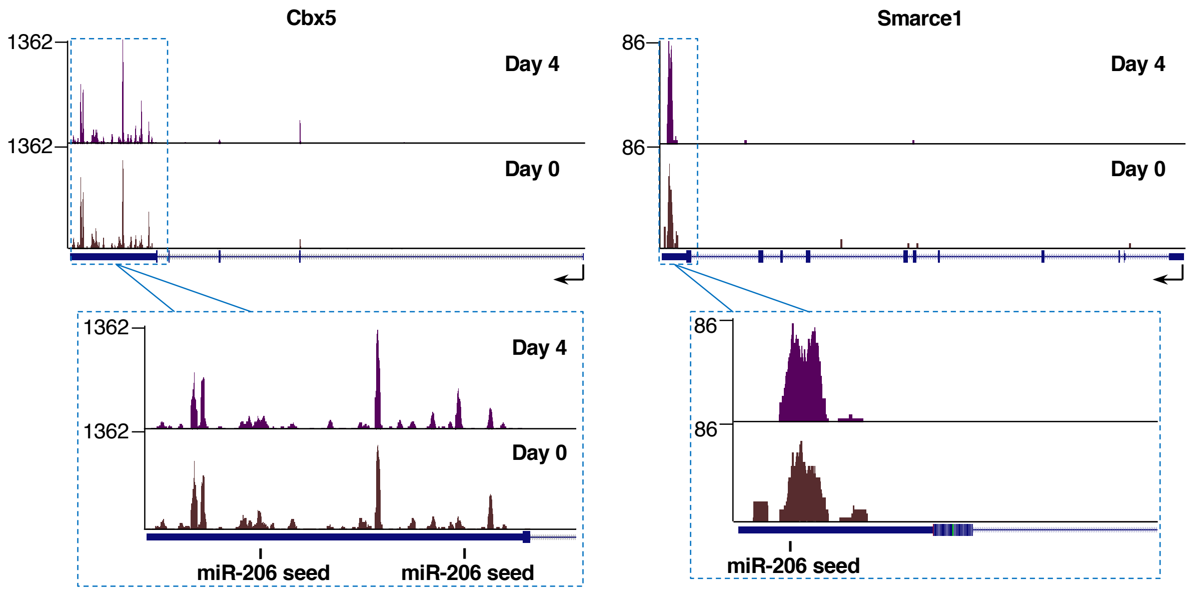


**Supplemental Figure 2.** xOP sequencing tracks for Cbx5 and Smarce1 show mapped reads enriched in the 3'UTRs of the genes. Reads pile up to form peaks over or near miR-206 seed sequences in the 3'UTRs of these genes. Sequencing tracks were created from the UCSC Genome Browser display of bedgraph files from replicate 1 mapped reads.

| Gene name | Replicate 1 Day 4 | Replicate 1 Day 0 | Replicate 2 Day 4 | Replicate 2 Day 0 |
|---|---|---|---|---|
| A330074K22Rik | 4.9 | 1.3 | 4.7 | 0.0 |
| Abraxas2 | 57.5 | 17.8 | 68.6 | 29.6 |
| Adam19 | 91.4 | 36.4 | 117.9 | 38.8 |
| Adamts2 | 19.8 | 7.3 | 17.6 | 2.9 |
| Adamtsl5 | 14.7 | 1.9 | 10.6 | 0.0 |
| Adcy9 | 10.8 | 1.7 | 8.0 | 2.3 |
| Adgrg6 | 12.3 | 4.6 | 16.8 | 2.4 |
| Agap1 | 20.0 | 4.1 | 22.5 | 9.0 |
| Apc | 10.4 | 3.6 | 13.3 | 3.8 |
| Arhgap23 | 9.8 | 2.0 | 7.5 | 0.0 |
| Arhgap28 | 10.8 | 1.7 | 11.1 | 0.0 |
| Arhgap28 | 10.7 | 1.7 | 11.1 | 0.0 |
| Asb1 | 27.1 | 12.7 | 31.6 | 13.7 |
| Atf5 | 296.1 | 127.6 | 519.8 | 155.3 |
| Atf5 | 168.0 | 74.0 | 269.4 | 88.7 |
| Atf6 | 90.4 | 25.4 | 113.3 | 30.1 |
| AU040320 | 10.2 | 0.0 | 14.6 | 3.5 |
| Bgn | 451.8 | 189.3 | 522.1 | 176.0 |
| C1qtnf3 | 40.3 | 8.7 | 61.3 | 4.6 |
| Camk2a | 109.7 | 1.5 | 123.8 | 2.0 |
| Camsap1 | 18.2 | 6.3 | 11.6 | 3.5 |
| Ccbe1 | 3.9 | 0.6 | 7.6 | 3.1 |
| Ccl9 | 12.2 | 4.2 | 11.7 | 1.4 |
| Cdkn1a | 564.3 | 101.7 | 758.5 | 48.6 |
| Cds2 | 40.7 | 15.8 | 36.9 | 15.1 |
| Cdyl2 | 12.9 | 5.7 | 18.7 | 3.7 |
| Chid1 | 5.2 | 0.0 | 8.2 | 2.3 |
| Chrna1 | 71.7 | 15.3 | 76.3 | 14.1 |
| Cln5 | 34.9 | 16.1 | 37.2 | 15.8 |
| Cnnm4 | 9.2 | 1.2 | 6.6 | 1.6 |
| Cnp | 10.6 | 0.0 | 15.3 | 3.6 |
| Cpeb4 | 64.6 | 24.7 | 83.1 | 16.6 |
| Csdc2 | 10.5 | 0.0 | 17.2 | 0.0 |
| Daam2 | 63.9 | 10.4 | 67.1 | 12.2 |
| Dclk1 | 21.6 | 2.4 | 14.4 | 1.3 |
| Dclk1 | 21.1 | 2.4 | 14.1 | 1.2 |
| Dennd5b | 19.1 | 5.1 | 11.9 | 4.8 |
| Dgkd | 110.3 | 21.9 | 130.9 | 33.7 |
| Dip2c | 20.7 | 9.0 | 29.8 | 4.7 |
| Dnal1 | 28.3 | 13.2 | 36.6 | 10.0 |
| Dnmt3a | 20.2 | 7.8 | 11.5 | 5.1 |
| Drp2 | 23.9 | 11.4 | 29.7 | 11.4 |
| Emp2 | 116.6 | 56.3 | 125.4 | 48.2 |
| Enah | 40.4 | 12.9 | 48.6 | 14.0 |
| Endov | 12.4 | 3.2 | 9.5 | 3.4 |
| Ezh1 | 41.1 | 10.0 | 42.1 | 16.8 |
| Fam53b | 10.5 | 1.5 | 10.8 | 0.0 |
| Fbn1 | 194.1 | 86.5 | 212.9 | 65.1 |
| Fbxw7 | 15.1 | 1.6 | 11.6 | 4.1 |
| Fem1a | 30.1 | 11.5 | 32.4 | 9.3 |
| Foxj2 | 9.9 | 2.1 | 11.5 | 2.7 |
| Galnt6 | 21.8 | 0.0 | 29.3 | 1.0 |
| Ghitm | 206.2 | 88.9 | 229.1 | 74.1 |
| Gpd1l | 48.9 | 18.7 | 40.7 | 16.6 |
| Grpel2 | 17.2 | 5.5 | 15.8 | 7.1 |
| Gstm2 | 28.8 | 0.0 | 27.7 | 9.8 |
| Gys1 | 84.0 | 18.7 | 61.4 | 13.6 |
| H19 | 261.8 | 96.7 | 320.5 | 74.8 |
| Hdac4 | 11.5 | 4.0 | 15.8 | 5.2 |
| Heyl | 18.5 | 5.6 | 22.2 | 5.2 |
| Hif1an | 63.5 | 23.4 | 65.7 | 20.6 |
| Hivep2 | 12.2 | 1.6 | 17.5 | 2.1 |
| Igf2 | 272.3 | 26.4 | 341.7 | 12.3 |
| Igfbp5 | 368.7 | 109.7 | 435.3 | 85.9 |
| Jph2 | 133.7 | 32.4 | 151.2 | 18.2 |
| Klhl31 | 56.3 | 4.3 | 73.2 | 6.4 |
| Ksr1 | 23.7 | 9.5 | 29.7 | 9.9 |
| Limd1 | 61.7 | 27.0 | 95.6 | 28.0 |
| Lrig2 | 15.2 | 6.9 | 16.5 | 7.2 |

| Gene name | Replicate 1 Day 4 | Replicate 1 Day 0 | Replicate 2 Day 4 | Replicate 2 Day 0 |
|---|---|---|---|---|
| Lynx1 | 33.3 | 9.2 | 59.7 | 3.0 |
| Map1a | 52.8 | 18.0 | 57.7 | 14.9 |
| Mapkbp1 | 15.7 | 2.5 | 11.6 | 4.9 |
| Marc2 | 64.3 | 28.7 | 52.9 | 16.7 |
| Mef2d | 61.6 | 14.3 | 69.6 | 20.7 |
| Mical3 | 8.8 | 0.8 | 13.8 | 1.1 |
| Micu2 | 68.3 | 31.6 | 121.5 | 20.6 |
| Mllt11 | 42.7 | 14.3 | 55.6 | 14.5 |
| Mlst8 | 7.7 | 1.1 | 6.3 | 1.5 |
| Mtf1 | 22.9 | 9.1 | 36.1 | 7.5 |
| Mylk4 | 59.2 | 8.3 | 79.8 | 6.2 |
| Ndrg1 | 92.0 | 23.5 | 83.7 | 28.5 |
| Nrbp2 | 8.4 | 3.5 | 9.7 | 2.3 |
| Nsmce2 | 18.7 | 7.3 | 20.2 | 9.5 |
| Nuak1 | 70.0 | 18.7 | 77.1 | 26.7 |
| Patl1 | 5.7 | 1.5 | 5.5 | 0.0 |
| Pbxip1 | 64.0 | 25.0 | 72.5 | 17.1 |
| Pcnx | 8.4 | 3.3 | 9.1 | 1.4 |
| Pde7a | 23.7 | 10.9 | 31.8 | 11.3 |
| Phf8 | 6.2 | 1.1 | 9.9 | 2.8 |
| Pkp1 | 147.6 | 36.9 | 130.0 | 27.2 |
| Pkp1 | 147.4 | 36.9 | 129.8 | 27.1 |
| Pmaip1 | 6.9 | 2.4 | 24.2 | 4.7 |
| Pnmal2 | 45.7 | 7.4 | 35.7 | 7.8 |
| Popdc3 | 79.8 | 27.7 | 112.3 | 36.2 |
| Prkaa2 | 16.7 | 3.2 | 23.8 | 2.6 |
| Pvr | 6.5 | 1.7 | 12.4 | 2.2 |
| Qsox1 | 81.4 | 38.5 | 99.4 | 33.5 |
| Rab5b | 14.9 | 6.6 | 12.3 | 4.3 |
| Rassf2 | 17.7 | 4.4 | 19.0 | 2.9 |
| Rgma | 6.7 | 0.0 | 15.4 | 0.0 |
| Rgp1 | 23.8 | 11.2 | 24.0 | 10.8 |
| Sema6a | 16.0 | 5.8 | 23.0 | 3.3 |
| Serpinb1a | 27.5 | 0.0 | 33.1 | 0.0 |
| Setbp1 | 7.2 | 2.7 | 8.9 | 2.8 |
| Sft2d2 | 50.9 | 21.7 | 57.6 | 12.5 |
| Sh3pxd2a | 88.1 | 42.7 | 85.2 | 30.9 |
| Sidt2 | 16.1 | 7.4 | 20.6 | 9.7 |
| Slc16a2 | 8.6 | 1.1 | 12.3 | 0.0 |
| Slc39a3 | 11.4 | 3.2 | 13.9 | 0.0 |
| Slc7a5 | 89.8 | 19.2 | 55.8 | 18.0 |
| Sort1 | 24.0 | 0.6 | 19.1 | 0.8 |
| Spg20 | 124.1 | 58.2 | 168.5 | 55.6 |
| Srl | 337.2 | 20.8 | 416.7 | 11.8 |
| Srl | 337.4 | 31.3 | 419.1 | 9.1 |
| Srr | 37.5 | 16.0 | 48.7 | 12.0 |
| Stard5 | 9.9 | 1.5 | 5.5 | 0.0 |
| Stbd1 | 261.0 | 114.3 | 290.1 | 93.4 |
| Stk25 | 13.0 | 3.0 | 5.5 | 2.0 |
| Stom | 28.1 | 11.1 | 41.0 | 3.6 |
| Stx17 | 21.0 | 9.8 | 29.2 | 8.6 |
| Susd6 | 133.4 | 47.6 | 143.5 | 30.2 |
| Tceanc2 | 8.3 | 1.0 | 6.2 | 0.0 |
| Tex2 | 15.4 | 7.1 | 19.8 | 4.7 |
| Tmcc3 | 33.1 | 9.3 | 48.9 | 10.4 |
| Tnfrsf23 | 11.9 | 0.0 | 7.0 | 1.2 |
| Tnfrsf23 | 11.8 | 0.0 | 7.0 | 1.2 |
| Tnfrsf23 | 11.7 | 0.0 | 6.9 | 1.2 |
| Tom1l2 | 13.3 | 6.2 | 14.2 | 6.0 |
| Top1 | 28.9 | 12.9 | 55.6 | 25.3 |
| Tpm2 | 142.4 | 30.8 | 148.7 | 23.2 |
| Trp53inp2 | 172.7 | 56.2 | 209.0 | 43.4 |
| Ttc9 | 27.1 | 7.9 | 26.7 | 6.9 |
| Usp20 | 10.7 | 4.4 | 12.3 | 2.9 |
| Zfp106 | 94.2 | 41.4 | 113.2 | 31.2 |
| Zfp322a | 20.1 | 8.0 | 17.8 | 6.3 |
| Zfp36l1 | 24.6 | 10.0 | 18.4 | 7.4 |
| 2010111I01Rik | 8.7 | 1.9 | 7.2 | 1.7 |
| 2010111I01Rik | 18.2 | 7.1 | 21.8 | 0.0 |

**Supplemental Table 1. Putative miR-206 targets identified through xOP-seq.** Shown are the normalized reads per kilobase in the 3'UTR for the 130 putative miR-206 targets identified via xOP-seq. Data for each replicate at Day 0 and Day 4 time points are shown.

| TERM | GENE COUNT | % | P-value | Benjamini |
|---|---|---|---|---|
| intracellular signaling transduction | 35 | 26.9 | 6.8E-6 | 1.8E-2 |
| signal transduction | 60 | 46.2 | 4.2E-5 | 5.5E-2 |
| signaling | 61 | 46.9 | 1.8E-4 | 9.0E-2 |
| single organism signaling | 61 | 46.9 | 1.5E-4 | 9.4E-2 |
| regulation of response to stimulus | 39 | 30.0 | 2.6E-4 | 9.5E-2 |
| cell communication | 61 | 46.9 | 2.4E-4 | 1.0E-1 |
| regulation of cellular protein metabolic process | 30 | 23.1 | 3.6E-4 | 1.0E-1 |
| regulation of molecular function | 29 | 22.3 | 4.1E-4 | 1.0E-1 |

**Supplemental Table 2. Eight biological pathways were enriched for the list of 130 putative miR-206 targets identified by xOP-Seq.** Biological pathways (category GOTERM_BP_ALL) were identified by DAVID's Functional Annotation tool. Enriched terms had Benjamini-Hochberg corrected *p* values $\leq 0.1$. Term refers to the enriched biological pathway. Gene count refers to the number of genes out of 130 in each term. Percent (%) refers to the percent of genes out of 130 in term. P-value refers to the uncorrected *p* values for each enriched term. Benjamini refers to the Benjamini-Hochberg corrected *p* values for each term.

| Gene name | Forward sequence | Reverse sequence |
|---|---|---|
| Adam19 | ACATGACCAGGATGCCACCAAACG | CACACCTCCAAGCCGACAAGTGC |
| Bgn | ATGACTTCAAAGGCCTCCAGCACC | AGTTTTTGCAGCTTCCGCAGAGGGC |
| Cbx5 | TGTTGGACAGGCGCATGGTTAAGG | GACAATCCAAGTTCTTCTCAGGTTCCC |
| Ccnd1 | GTGCAGAAGGAGATTGTGCCATCC | GCGGTCCAGGTAGTTCATGGC |
| Csnk2a2 | GAGCTTGGGCTGCATGTTAGCG | GTTCATCTGTCCCCAGAACCTTGGC |
| Gja1 | GTCAACGTGGAGATGCACCTGAAGC | TGCTGATGATGTAGGTTCTCAGCAGG |
| Hdac4 | CAATGCCAATGCTGTCCACTCCATGG | CTTTTGCGCCTCAATCAGAGAGTGC |
| Smarce1 | GAAGCCGAGCTCCTTCAGATAGAGG | GATCTCAGCCGCAATCTTCTCCATG |
| Spg20 | CTCAACTGTCTGGCAAGGGTTGG | GTAGCTTCTCCTGCGTTGTGCC |
| Tmcc3 | CAGGCCTATGAGCGCTCAAGG | TTCACGGCATCCGTCTGCAGG |

**Supplemental Table 3. qPCR Primers.** Sequences are written 5’ to 3’.
